# Intracellular *Plasmodium* aquaporin 2 is required for sporozoite production in the mosquito vector and malaria transmission

**DOI:** 10.1101/2023.03.15.532816

**Authors:** Alexander J. Bailey, Chiamaka Valerie Ukegbu, Maria Giorgalli, Tanguy Rene Balthazar Besson, George K. Christophides, Dina Vlachou

## Abstract

Malaria remains a devastating disease and, with current measures failing to control its transmission, there is a need for novel interventions. A family of proteins that have long been pursued as potential intervention targets are aquaporins which are channels facilitating the movement of water and other solutes across membranes. We identify a new aquaporin in malaria parasites and demonstrate that it is essential for disease transmission through mosquitoes. Disruption of AQP2 in the human parasite *Plasmodium falciparum* and the rodent parasite *Plasmodium berghei* blocks sporozoite production inside oocysts established on mosquito midguts, preventing parasite infection of salivary glands and transmission to a new host. *In vivo* epitope tagging of AQP2 in *P. berghei*, combined with immunofluorescence assays, reveals that the protein is localized in previously uncharacterized organelles found in the cytoplasm of gametocytes, ookinetes and sporozoites. The number of these organelles varies between individual parasites and lifecycle stages suggesting that they are likely part of a dynamic endolysosomal system. Phylogenetic analysis confirms that AQP2 is unique to malaria and closely related parasites and most closely related to other intracellular aquaporins. Structure prediction analyses identify several unusual features, including a large accessory extracellular loop and an arginine-to-phenylalanine substitution in the selectivity filter principally determining pore function, a unique feature not found in any aquaporins studied to date. This in conjunction with the requirement of AQP2 for malaria transmission suggests that AQP2 may be a fruitful new target of novel antimalarial interventions.

## Introduction

Malaria, caused by protozoan parasites of the *Plasmodium* genus, remains a significant global health challenge despite recent successes against the disease. In recent decades, progress towards control of malaria has stalled, partly driven by growing parasite resistance to treatment, fueled by rising mosquito resistance to insecticides used in bed nets and indoor residual spraying. Historically, targeting the mosquito vector has yielded great success in controlling malaria cases, and has led to disease eradication in some regions. Given the diminishing impact of established malaria interventions, it is of paramount importance to identify new targets for development of novel malaria interventions to control the disease. Here, we describe the identification and characterization of one such putative target: a new aquaporin, AQP2, which we show to be essential for completion of *Plasmodium* development within the mosquito vector and production of sporozoites, the parasite form responsible for transmission to humans.

Aquaporins are transmembrane proteins that facilitate the movement of water and other solutes across membranes. They are broadly grouped into exclusive water transporters (orthodox), water channels with solute permeability (*e.g*., aquaglyceroporins that transport glycerol) and less well-studied intracellular aquaporins (unorthodox or super-aquaporins), although this classification belittles the diversity of aquaporins in terms of both localization and selectivity (1). Aquaporins share a common tertiary structure, with six transmembrane helices arranged in an hourglass pore-forming shape, joined by five extramembrane loops. Two half-helices that dip into either side of the membrane and face each other in the pore center contribute to the pore selectivity and function. Interestingly, aquaporins form tetramers creating a central pore that may additionally transport solutes including ions and small molecules (2–6). Whilst aquaporins on the plasma membrane facilitate exchange between the cell and its environment, intracellular aquaporins facilitate exchange between cytoplasm and organelles. For example, human AQP11 transports H_2_O_2_ into and out of the ER (7), which may explain its essential role in the kidney (8), while AQP1 of the apicomplexan parasite Toxoplasma gondii is localized to a plant vacuole-like organelle that serves as a sodium/calcium store and confers stress resistance (9).

*Plasmodium* AQP1 is a well-characterized plasma membrane aquaglyceroporin that conducts water, glycerol, and ammonia at high rates (10–12). It is expressed throughout parasite asexual blood stages (ABS), sporozoites and infected hepatocytes, and provides parasites access to serum glycerol for membrane lipid synthesis (10, 11, 13). Whilst the essentiality of AQP1 to ABS is disputed (11, 14), it is established that the protein is important for progression of hepatic stage infection (13). Interestingly, amino acid residues contributing to AQP1 pore selectivity are identical with those in E. coli glycerol facilitator GlpF, which in conjunction with high conservation between the two proteins suggests that AQP1 has been acquired from a prokaryotic ancestor (10, 15).

A second putative aquaporin, AQP2, has been identified in *Plasmodium* genomes and reported to be phylogenetically associated more closely to *T. gondii* AQP1, rather than *Plasmodium* AQP1 (14). We previously performed a reverse genetic screen in the rodent model parasite *P. berghei* of genes with enriched expression in gametocytes, which identified AQP2 as important for mosquito salivary gland infection (16). Here, we aimed to follow up on these results and characterize AQP2 in both *P. berghei* and the human parasite *P. falciparum*. Generation and phenotypic characterization of knockout parasites confirmed that AQP2 is essential for production of sporozoites. Immunofluorescence assays of tagged AQP2 showed that the protein is intracellular, residing in a vesicle-like organelle of female gametocytes, ookinetes and sporozoites. Phylogenetic analysis along Alphafold structure predictions revealed that AQP2 is distinct from AQP1, other aquaglyceroporins and orthodox aquaporins with regards to both its overall architecture and pore selectivity filter. Our results validate AQP2 as a potential transmission blocking target of future antimalarial interventions.

## Results

### AQP2 is distinct from AQP1

AQP2 orthologs are found in all plasmodia with available genome sequences. They encode large aquaporin-like proteins of 603 amino acids in *P. falciparum* (PfAQP2, PF3D7_0810400) and 595 amino acids in *P. berghei* (PbAQP2, PBANKA_1427100). Multiple sequence alignment identified regions of high conservation mostly associated with predicted a-helixes (Fig. S1). One region is substantially divergent between species, both in terms of sequence identity and length: amino acids 110-228 in PfAQP2 and 110-215 in PbAQP2. Insertions in this region cause AQP2s of human parasites of the sub-genus *Plasmodium* (*P. malariae, P. vivax* and *P. knowlesi*) to be considerably longer than other AQP2s. Protein structure predictions made with Alphafold Monomer v2.0 (17, 18) showed that PfAQP2 and PbAQP2, like all aquaporins, have six full transmembrane helices and two half helices, connected by five loops (Fig. S2). The variable region corresponds to extracellular loop A that lies between helixes 1 and 2. Loop A is highly charged, with roughly 40% of amino acids carrying either a positive or negative charge. Intracellular loop D is longer than extracellular loop C, the inverse of loop lengths of *Plasmodium* AQP1. The characteristic aquaporin pore forming NPA motifs are present in all AQP2s, in half-helices 1 and 2 that are part of loops B and E, respectively. While the first NPA motif is present as a canonical NPA, the second motif is found as NPM in all species except *P. gallinaceum* which harbors an A/M>L substitution. Finally, in all AQP2s, the canonical arginine residue that determines the selectivity filter, invariably found directly adjacent to the second NPA motif, is replaced with phenylalanine (R>F).

To understand the evolutionary history of *Plasmodium* AQP2s, we performed a phylogenetic analysis of the conserved pore-forming regions of AQP2s and 66 other aquaporins from 21 other species across all domains of life, including *Homo sapiens* and *Anopheles gambiae* – the vertebrate and mosquito hosts of *P. falciparum*, respectively (Fig. S3). A clear clustering of aquaporins was observed principally by the type of pore formed, whether orthodox, solute transporters including aquaglyceroporins, or unorthodox. *Plasmodium* AQP2s form a well-defined clade together with an aquaporin of the closely related parasite *Hepatocystis*, which appears to be related to intracellular *T. gondii* AQP1 (TgAQP1). A clade involving *A. gambiae* AQP7 and the unorthodox human aquaporins AQP11 and AQP12 are the closest to *Plasmodium* AQP2s and TgAQP1, although this relation lies deep in the root of the tree and is not supported by bootstrapping. *Plasmodium* AQP1s form a separate clade together with human aquaglyceroporins and *E. coli* glycerol facilitator GlpF, as previously described (10).

From these data, we concluded that AQP2 most closely resembles intracellular, unorthodox aquaporins, with their pore-forming sequences closely related to *Hepatocystis* aquaporin and, albeit less so, to the intracellular, vacuole-localized TgAQP1. The arginine to phenylalanine substitution in the AQP2 selectivity filter is unique to *Plasmodium* and, to our knowledge, not found in any other aquaporin.

### Alphafold structure predictions reveal unusual AQP2 pore and selectivity filter features

We used Alphafold Monomer v2.0 (17, 18) to study the putative three-dimensional structure of PfAQP2 and PbAQP2 (UniProt C0H4T7 and A0A509AUY5, respectively). The results showed that the highly conserved pore-forming helices have the highest confidence structure predictions (Fig. S4). However, helix 6 alongside loop E of PbAQP2 has low confidence prediction and does not correctly fold into the core as expected. Therefore, we proceeded with analysis of the PfAQP2 structure as absence of helix 6 from the PbAQP2 structure may result in unintended changes to the pore. Nevertheless, all the observations below match those also made for PbAQP2 (Fig. S5).

Overall, the Alphafold PfAQP2 structure provides high confidence predictions of the pore-forming regions including the selectivity filter and downstream NPA/NPM motifs (Fig. 1*A*). The four residues constituting the constriction at the selectivity filter are: Ile62 (helix 1), Phe465 (helix 5), Val468 (helix 5) and Phe477 (half helix 2). Whilst Phe477 on the site canonically occupied by an arginine residue does not appear to affect the formation of the pore, the presence of two aromatic phenylalanine residues opposite each other is a major difference from orthodox aquaporins and may pose a restriction to the size of the pore. The isoleucine residue is contributed to the pore by helix 1 as opposed to helix 2 as in most other aquaporins, causing a substantial tilt of AQP2 in the membrane axis, although this does not change the predicted structure of the pore. The NPA and NPM motifs contributed by half-helices 1 and 2, respectively, meet one another in the pore. Interestingly, the central pore is mostly nonpolar; however, a positively charged histidine residue (His302) found near the interior side of the NPA/NPM motif places a positively charged residue close to the pore center.

**Fig. 1.**
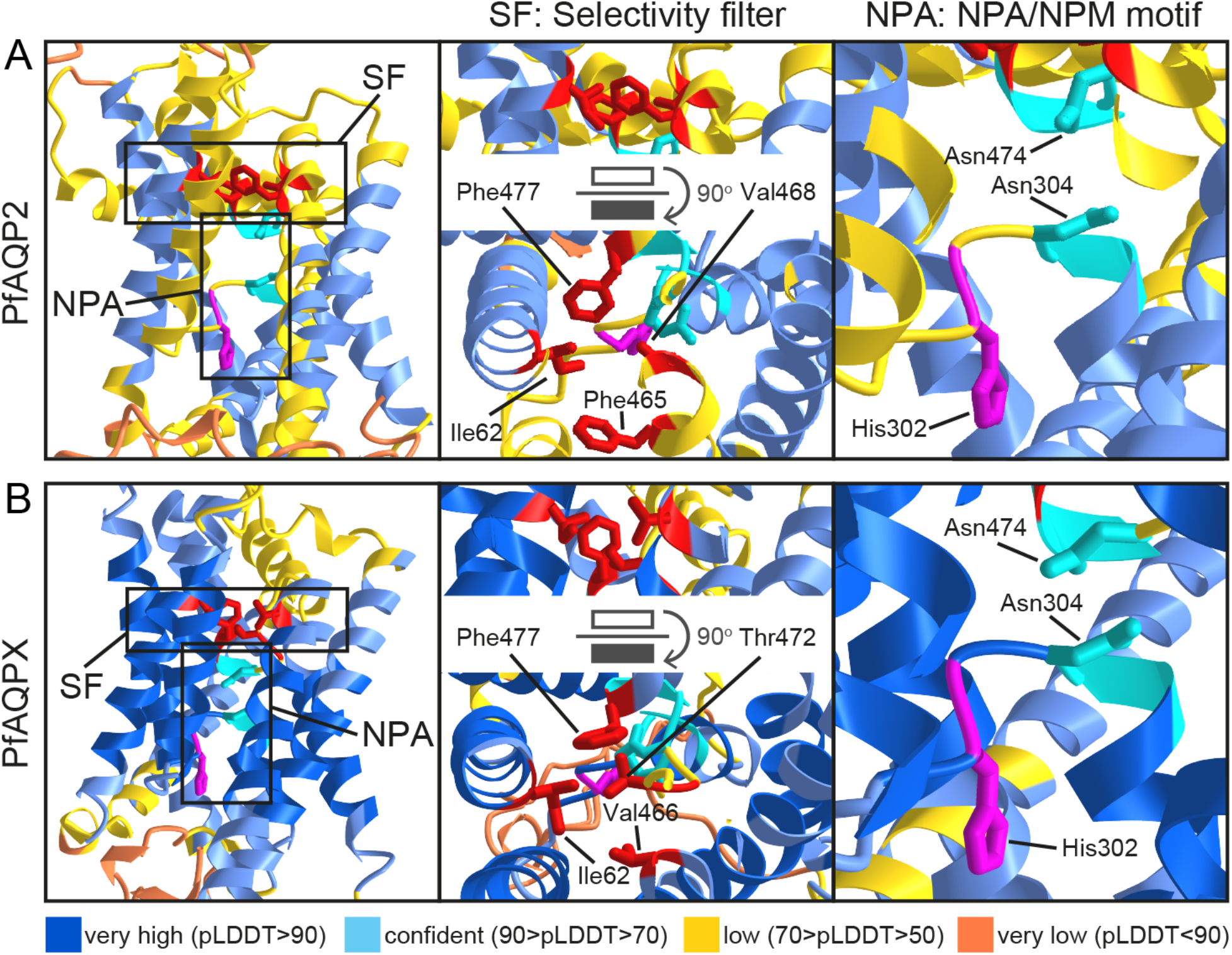
Predicted structure of PfAQP2 pore and selectivity filter. (*A*) PfAQP2. (*B*) Chimeric PfAQPX in which PfAQP2 loops A-E are replaced with corresponding loops of previously resolved PfAQP1 structure. Structures are predicted with Alphafold Monomer v2.0 on the CoLab server. Top ranking models are shown. Structures are colored by confidence level as shown in the key below the figure. Residues contributing to the selectivity filter are shown in red and those contributing to the NPA/NPM motif are shown in cyan, while the histidine residue participating in the pore is shown in magenta.

To generate more high-confidence pore predictions, we generated a chimeric sequence with extramembrane PfAQP2 loops A-E replaced with the shorter loops of PfAQP1 that has a resolved tertiary structure (15). The chimeric protein is referred to as PfAQPX. Folding predictions using Alphafold on the CoLab server indeed generated much higher confidence predictions of PfAQPX pore structure (Fig. 1*B*). Whilst most PfAQP2 pore features and structure were preserved in PfAQPX, Phe465 was rotated away to no longer participate in the filter and instead replaced by Thr472 (half helix 2). All the rest of the amino acid residues remained the same as in the PfAQP2 structure, including Phe477.

### PfAQP2 and PbAQP2 are essential for sporozoite maturation in the oocyst

Our earlier work has shown that PbAQP2 expression is higher in gametocytes compared to ABS (16), and this is confirmed by single cell transcriptome analysis in the Malaria Cell Atlas (19). We sought to study the expression of AQP2 in *P. falciparum* by quantitative real-time RT-PCR (qRT-PCR). In ABS, there is very low expression of AQP2, in contrast to *in vitro* cultured gametocytes that express the highest amount of AQP2 in all stages assayed (Fig. 2*A*). In *A. coluzzii* infections, highest AQP2 expression was detected in the blood bolus 1hour pbf (post blood feeding), a signal that most likely derives from gametocytes, which remains high 24 hours pbf, a timepoint that also encompasses mature ookinetes. Following establishment of oocysts, AQP2 transcription appeared to be minimal 6 days pbf, but a slight increase was observed 9 days pbf, possibly associated with re-expression in maturing sporozoites.

**Fig. 2.**
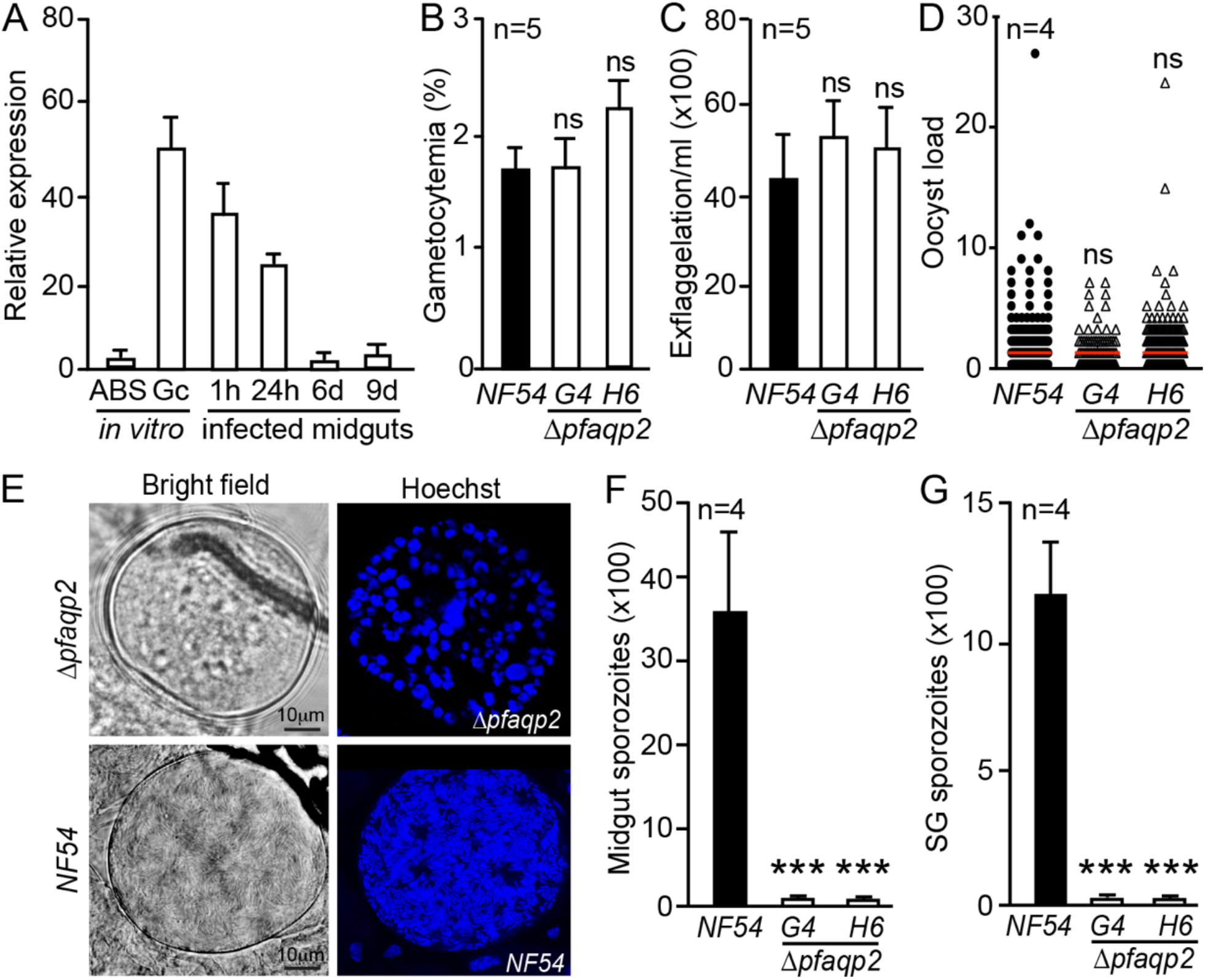
Phenotypic characterization of *P. falciparum* AQP2. (*A*) QRT-PCR of AQP2 transcripts in *in vitro* cultured asexual blood stages (ABS) and mature gametocytes (Gc) and in *in vivo* infections of *A. coluzzii* midguts at 1 hour (1h), 24 hours (24h), 6 days (6d) and 9 days (9d) post blood feeding (pbf). Relative expression was calculated for each sample relative to *in vitro* and *in vivo* internal calibrator reference values using EF1α as a housekeeping gene. Mean and SEM values from 3 independent replicates are presented. (*B*) Percentage gametocytemia in total parasite numbers in *in vitro* gametocyte culture. Mean and SEM values from 5 independent replicates are shown. (*C*) Number of exflagellation centers per 1 mL of *in vitro* gametocyte culture. Mean and SEM are shown. (*D*) Oocyst load in mosquito midguts 9 days pbf. Medians from pools of 4 replicates are presented (red lines). (*E*) Bright field and Hoechst imaging of oocysts 11 days pbf. Scale bar is 10 μm. (*F*) Midgut and (*G*) salivary gland sporozoites enumerated using a hemocytometer with mean and SEM presented. In all panels, results from *NF54* control and two independent *Δpfaqp2* parasite lines (G4 and H6) are presented. Statistical analysis is done with ANOVA for data in panels B-C and F-G and Kruskal-Wallis test for data in panel D. ns means not significant and *** shows *p*< 0.001.

We generated a PfAQP2 deletion mutant via CRISPR/Cas9 and single homologous recombination (Fig. S6A). Successful integration of a selectable marker in *P. falciparum* NF54 and limiting dilution cloning was confirmed by diagnostic PCR and led to the generation of two successful *Δpfaqp2* clones (G4 and H6; Fig. S6*B*). The capacity of *Δpfaqp2* to form gametocytes was assessed 14 days after induction of *in vitro* cultured ABS. No significant difference was observed between NF54 and the *Δpfaqp2* lines in the establishment of mature stage V gametocytes (Fig. 2*B*). Next, we performed *in vitro* exflagellation assays to assess the capacity of *Δpfaqp2* to produce male gametes by counting exflagellation centers. Again, no difference was observed between the *NF54* control and the two *Δpfaqp2* lines (Fig. 2*C*).

Oocyst presence on the midguts of *A. coluzzii* 9 days pbf on *in vitro* gametocyte cultures was comparable between *NF54* control and *Δpfaqp2* lines, both in terms of load, *i.e*., number of oocysts per midgut, and prevalence, *i.e*., proportion of mosquitoes harboring oocysts (Fig. 2*D* and Table S1). However, imaging oocysts 11 days pbf suggested that nuclear division into daughter nuclei for budding sporozoites was less advanced in *Δpfaqp2* compared to *NF54* control oocysts (Fig. 2*E*). Most, albeit not all, *Δpfaqp2* oocysts exhibited larger nuclei and smoother appearance compared to *NF54* oocysts, lacking the characteristic convolutions associated with budding sporozoites. Indeed, mechanical disruption of oocysts followed by counting of sporozoite numbers using hemocytometer confirmed almost total absence of mature sporozoites from *Δpfaqp2* oocysts 11 days pbf compared to thousands of sporozoites per mosquito harboring *NF54* oocysts (Fig. 2*F* and Table S2). Next, we enumerated sporozoites in mosquito salivary glands 18 days pbf. The results corroborated the oocyst findings, as only very few *Δpfaqp2* compared to thousands of *NF54* sporozoites were detected per mosquito (Fig. 2*G* and Table S2). These data led us to conclude that loss of *P. falciparum* AQP2 results in a dramatic reduction in development of mature sporozoites, but has no impact on any of the preceding stages.

We extended our investigation to AQP2 of *P. berghei* as this parasite allows more detailed analysis and mosquito-to-mouse transmission experiments. *P. berghei* mutant lines carrying a disrupted version of AQP2 were generated using a *PlasmoGEM* vector that replaced 70% of the AQP2 coding sequence with the *human DHFR (hDHFR)/yFCU* pyrimethamine resistance cassette in the *c507* GFP-expressing wildtype line (20). Integration of the disruption cassette and gene deletion in clonal *Δpbaqp2* was confirmed by PCR (Fig. S7*A*).

Male gametogenesis, determined as number of exflagellation events per male gametocytes, was comparable between *Δpbaqp2* and the parental *c507* control line (Fig. 3*A*), as was the capacity of female *Δpbaqp2* gametocytes to form ookinetes (Fig. 3*B*). Next, we assessed the ability of *Δpbaqp2* to form oocysts following feeding of *A. coluzzii* mosquitoes on infected mice. Oocyst enumeration in mosquito midguts 8 days pbf showed no difference in oocyst load and prevalence between *Δpbaqp2* and *c507* control parasites (Fig. 3*C* and Table S3).

**Fig. 3.**
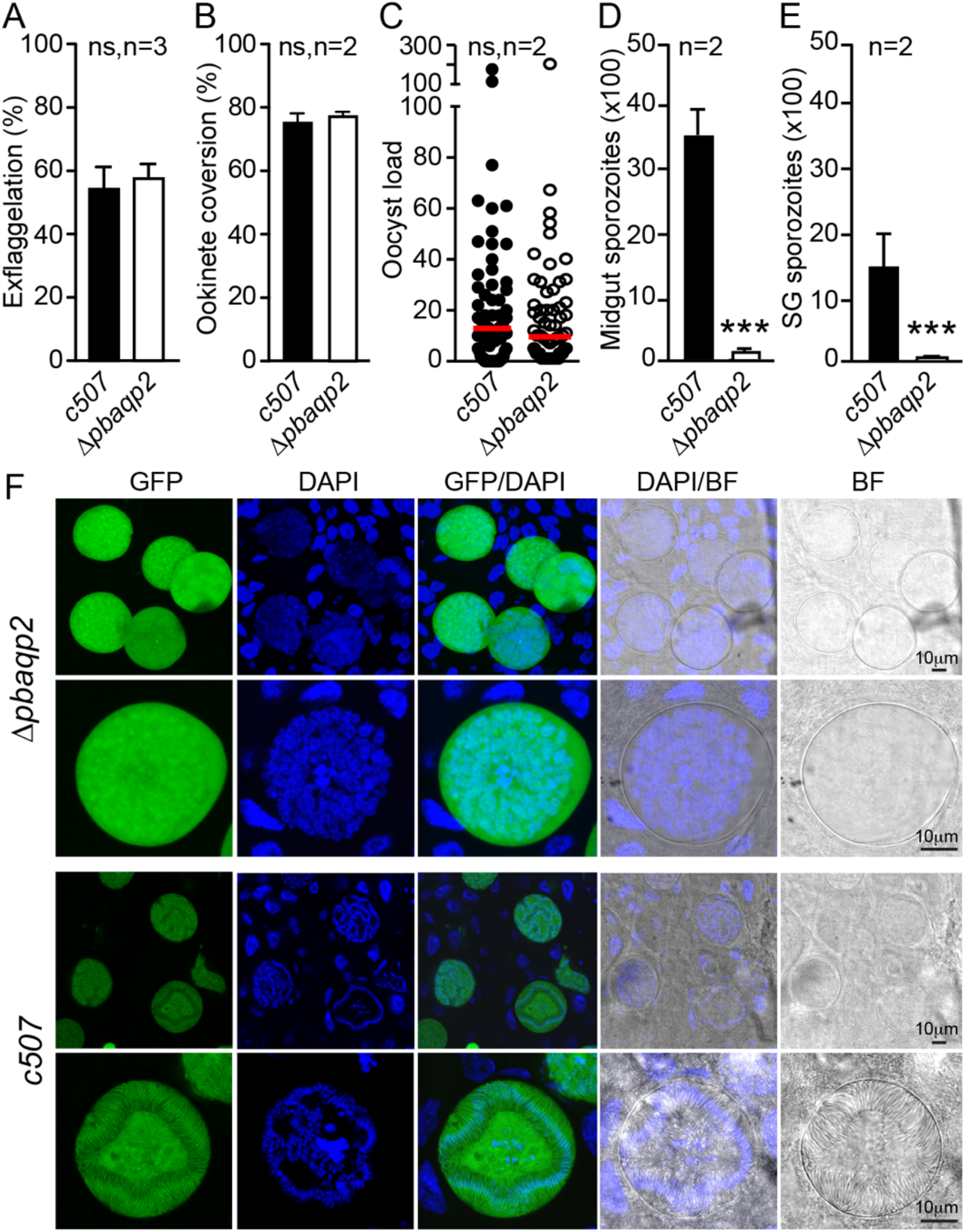
Phenotypic characterization of *P. berghei* AQP2. (*A*) Percentage *in vitro* exflagellation of male gametocytes. (B) Female gamete to ookinete conversion. (C) Oocyst load in the midguts of *A. coluzzii* mosquites enumerated 8 days pbf. Red lines indicate median. (D) Fluorescence microscopy images of GFP-expressing *Δpbaqp2* and *c507* control oocysts at day 15 pbf in *A. coluzzii* midguts. Bright field (BF) and DAPI staining are also shown. Scale bar is 10 μm. Graphs A, B, D and E depict mean and SEM with statistical analysis done with Student’s t-test. Graph C shows median and distribution with statistical analysis done with Mann-Whitney U test. n is number of independent biological replicates; ns is not significant; *** is *p*<0.001.

Next, we examined whether *Δpbaqp2* oocysts can produce sporozoites, firstly by counting sporozoites in mechanically ruptured oocysts 15 days pbf and secondly by counting sporozoites that had invaded the mosquito salivary glands 21 days pbf. The results fully matched those obtained with the *Δpfaqp2* parasites: both *Δpbaqp2* oocyst and salivary gland sporozoites were severely reduced compared to the *c507* controls (Fig. 3*D,F* and Table S4). DAPI staining of *Δpbaqp2* oocysts at day 15 pbf revealed that nuclei were large and disorganized unlike *c507* oocysts that had highly organized nuclei, characteristic of sporozoites that had budded off from the central sporoblastoid body (Fig. 1*E*). Finally, we tested whether *Δpbaqp2* parasites could be transmitted between vertebrate hosts through mosquito bites using bite-back assays on highly susceptible C57/BL6 mice 21 days pbf. Mice were monitored for infection daily for 14 days post mosquito bite. The almost total absence of *Δpbaqp2* sporozoites in the mosquito salivary glands led to termination of transmission.

### AQP2 resides in a new ookinete and sporozoite vesicle-like organelle

We tagged PbAQP2 at its C-terminus with a 3x human influenza hemagglutinin (3xHA) epitope via double crossover homologous recombination in the *c507* line to study protein expression. The new transgenic parasite line was named *aqp2::3xha* (Fig. S7*B*). Western blot analysis of *aqp2::3xha* parasite protein extracts using an antibody against the HA epitope confirmed that recombinant AQP2::3xHA is produced at the expected ca. 73 kDa size predominantly in gametocytes – already prior to gametogenesis activation – and, at a lower abundance, in mature ookinetes (Fig. 4*A*). Protein traces observed in mixed blood stages are thought to derive from gametocytes.

**Fig. 4.**
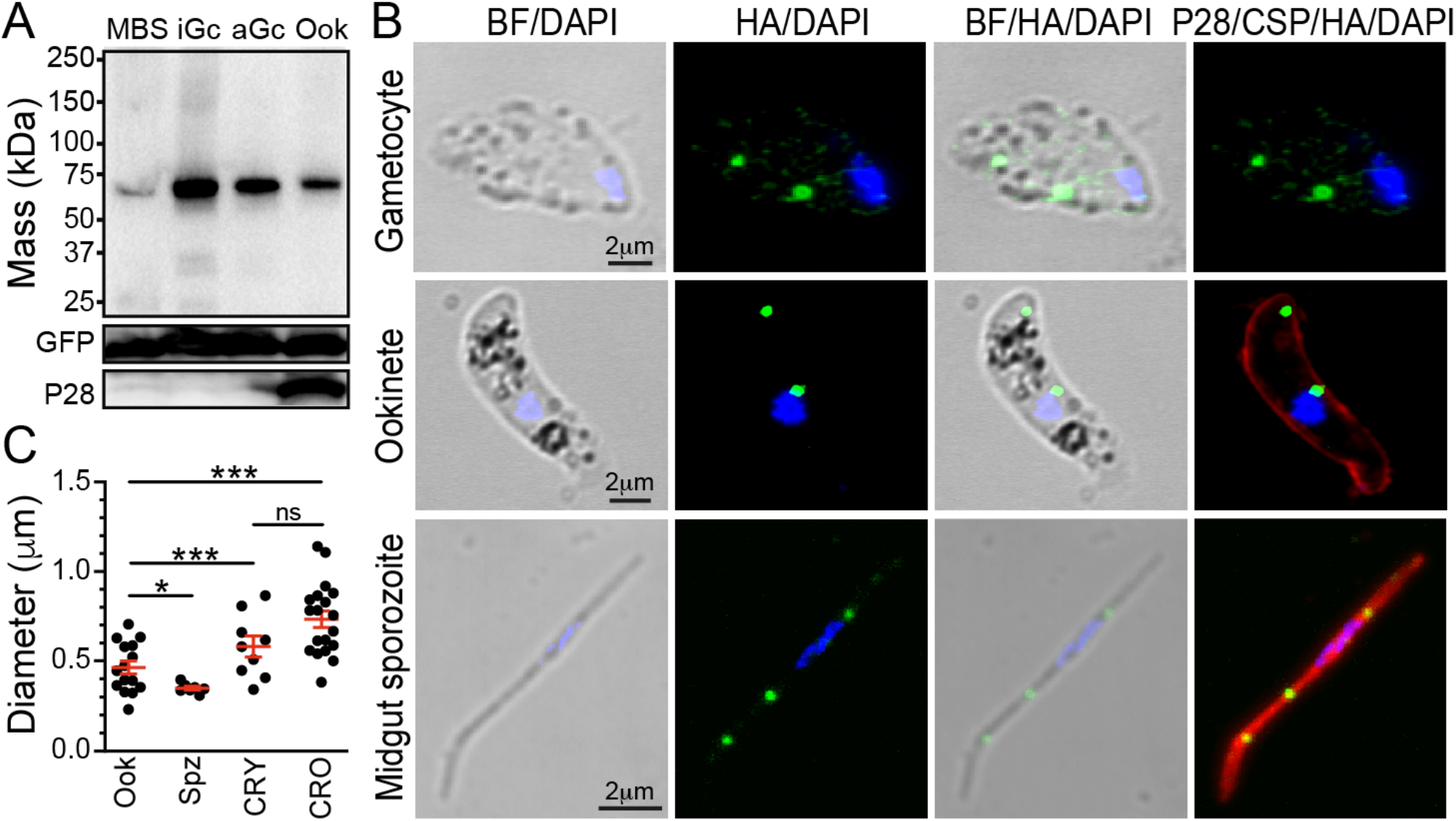
AQP2 protein expression and localization. (*A*) Western blot analysis on whole cell lysates from the *aqp2::3xha* parasite line under reducing conditions using α-HA antibody. GFP and P28 are used as loading and stage-specific controls, respectively. MBS is mixed blood stages, iGc is inactivated gametocytes, aGc is activated gametocytes and Ook is ookinetes. (*B*) Immunofluorescence assays of *aqp2::3xha* purified gametocytes, *in vitro* cultured ookinetes sampled 24 hours after culture setup, and sporozoites obtained from infected *A. coluzzii* midguts 9 days pbf. AQP2::3xHA (green) and P28 (red, ookinetes) or CSP (red, sporozoites) are detected using antibodies. DNA is stained with DAPI (blue). BF is brightfield. Scale bar is 2 μm. (*C*) Diameter of AQP2::3xHA stained vesicle-like structure in ookinetes (ook) and sporozoites (Spz), compared to diameter of crystalloids determined via CRYSP::3xHA (CRY) and CRONE::3HA (CRO) proteins.

In indirect immunofluorescence assays of *aqp2::3xha* parasites, AQP2::3xHA was seen to specifically localize to vesicle-like structures in the cytoplasm of gametocytes, ookinetes and midgut sporozoites (Fig. 4*B*). The number and appearance of these structures varied from 1-2 large spots and several smaller ones in gametocytes, 2-3 and occasionally 1-4 spots in ookinetes, and 2-3 spots in sporozoites (Fig. 4*B* and Fig. S8). In ookinetes, this pattern resembles that of crystalloid, an elusive organelle exclusive to ookinetes and young oocysts. Although these spots were often seen adjacent to the hemozoin crystals, the waste product of hemoglobin metabolism and hallmark of the ABS food vacuole and the crystalloid, they never fully colocalized with the hemozoin or located amongst the hemozoin crystals where the crystalloid is found. Indeed, the diameter of these structures in ookinetes (0.2-0.7 μm) was smaller than that of crystalloids (0.3-1.15 μm) as determined by IFAs of the recently identified crystalloid proteins CRYSP and CRONE (16) (Fig. 4*C*). The diameter of these structures was more uniform in all sporozoites examined (0.25-0.4 μm).

## Discussion

We demonstrate that *Plasmodium* AQP2 is an intracellular aquaporin residing in previously uncharacterized vesicle-like structures of gametocytes, ookinetes and sporozoites. Cellular bodies with “more regular shape and uniform structure than crystalloids”, resembling lysosomes, have been described already in 1962 by Garnham and coworkers (21). Whilst the crystalloid is now known to store proteins essential for oocyst development (22), these other cellular bodies have not been studied further. Given the appearance and size of the AQP2-inhabiting structures as well as their varying number between parasites and lifecycle stages, we postulate that these are part of the endolysosomal system, probably corresponding to the aforementioned bodies. Importantly, one of the closest phylogenetic relatives of AQP2, the intracellular *T. gondii* AQP1 resides in a multivesicular organelle presenting similarities to the plant vacuole, named plant-like vacuole (PLV) (9). Plants have two types of vacuoles: a lytic vacuole, which contains hydrolytic enzymes and functions as a digestive organelle similar to lysosomes, and a protein-storage vacuole (23). Intriguingly, plant vacuoles contain Tonoplast Intrinsic Proteins (TIP) aquaporins (24) that are related to *T. gondii* AQP1 (25). Therefore, we speculate that while the crystalloid is equivalent to the plant protein-storage vacuole, the AQP2-containing bodies are part of a multivesicular endolysosomal organelle equivalent to the lytic vacuole. In this organelle, AQP2 could play a role in water or energy storage or trafficking of essential solutes, similar to the roles observed in *T. gondii* AQP1 or human AQP11 (7–9, 25), serving an essential role in homeostasis, regulating the organelle pH, osmolarity or redox status. Interestingly, whilst both the crystalloid and the AQP2-containing vesicles are prominent in the ookinete, the essential function of their resident proteins is seen days later in the maturing oocyst.

The detection of AQP2-harboring vesicles in the sporozoite presents an additional challenge, as there is no reference of lysosomes in the sporozoite. A number of prominent vesicles in the region between the nucleus and apex, named micropores from their presumed relation to the parasite mouth-like structure, have been reported in early studies (26–29). Their increase in numbers late in sporozoite development made authors suggest an important function in parasite maturation. Micropore vesicles are up to 0.25 μm in diameter, which matches the size of AQP2-harboring vesicles (fluorescence measurements are expected to inflate vesicle size), and filled with granular material similar to those present in the oocyst matrix; therefore, they have been interpreted as endocytic vesicles. Interestingly, endocytic vesicles with similar contents have been described in *T. gondii* (30). Although these may be related to the PLV, no such connection has been made in the literature. Indeed, a later study has concluded that it is impossible to determine whether these vesicular structures are being trafficked in or out of the parasite (31).

Like *T. gondii* AQP1, *Plasmodium* AQP2 harbors an amino acid substitution at the critical arginine aromatic (Arg/R) region of the pore. Whilst in TgAQP1 arginine is replaced by a valine (25), a substitution characteristic of plant TIP-like aquaporins, in the case of AQP2 arginine is replaced by phenylalanine. Both amino acid residues are hydrophobic but phenylalanine is also aromatic and larger than valine. Phenylalanine is commonly found in orthodox aquaporin and aquaglyceroporin filters but not in the place of the canonical arginine, a feature unique to AQP2. In addition, the presence of two phenylalanine residues in the selectivity filter would make the AQP2 pore highly hydrophobic. It is possible, however, that this predicted structure is due to unconfident Alphafold predictions, stressing the need for future structural studies on purified AQP2. Indeed, when all the loops of AQP2 are replaced with those of AQP1, resolving a far more confident Alphafold prediction of the central pore region, the common phenylalanine residue in the selectivity filter is replaced by threonine, a smaller and polar residue, thus leaving only the unusually placed phenylalanine in the pore. Notably, whereas the replaced phenylalanine appears not to protrude directly into the pore, but rotated away from its central axis, threonine projects much further towards the pore center, resulting in a tighter constriction. Whether there is a phenylalanine or threonine in the AQP2 pore would most certainly alter solute selectivity by changing both the polarity and diameter of the pore. Nevertheless, both possible filters are mostly non-polar, and lack charged residues. Interestingly, in all predictions, isoleucine is contributed to the selectivity pore by helix 1 instead of helix 2 as in other aquaporins with resolved structures. This skews the structure of AQP2 to give it a substantial tilt in the membrane axis. Overall, *Plasmodium* AQP2 is almost certainly not an orthodox aquaporin.

In addition to its divergent pore, *Plasmodium* AQP2 harbors a number of interesting features in its extramembrane loop regions, especially extracellular loop A that is much larger in size than other aquaporins albeit variable between *Plasmodium* orthologs. This loop is typically short in known aquaporins; for example, in AQP1 it is only 6 amino acids long (10, 15). Examples of an extended loop A include in plant PIPs, where it appears to be involved in disulphide bonding of heterotetramers and can form a cap gating the central pore (32). In addition, in PIP2;1, the negative charges present in loop A are suggested to play a role in ion conduction through the pore. Although no cysteines are present in either PfAQP2 or PbAQP2 loop A, there is a high number of charged residues. For example, PfAQP2 loop A has 39/211 positively charged and 44/211 negatively charged residues. Therefore, loop A through its net-negative charge may regulate cation conductance through a putative AQP2 tetramer, like PIP2;1, be involved in gating of either the monomer or central pore, or facilitate interaction with other regulatory proteins, similar to *Saccharomyces cerevisiae* aquaglyceroporin FPS1. There, unusually large N and C-terminal extensions are used to regulate the channel permeability in response to stress through the recruitment of regulatory proteins such as kinase Hog1 and regulators of glycerol channel Rgc1 and Rgc2 (33, 34).

Whatever its molecular and cellular function, AQP2 is essential for sporozoite maturation and malaria transmission. This leads us to propose that AQP2 is a promising target of transmission-blocking drugs, given its prominent presence in blood-stage gametocytes, or other antimalarial interventions in the vector. There has been long and consistent interest in the pursuit of small molecules and biologics capable of blocking aquaporin channels. Although much of the focus to date has been on diseases such as cancer, metabolic disorders and oedema, the ubiquity, importance and divergence of aquaporins in protozoan parasites makes them potential targets for infectious disease control (35). Indeed, a recent study reports inhibition of *P. falciparum* growth through blocking AQP1 with the sweetener erythritol (36). The unique evolutionary history of *Plasmodium* AQP2 and divergence of its selectivity filter to known aquaporins, makes it a fruitful target of small molecules with minimal cross-reactivity to aquaporins of host and vector species.

## Materials and Methods

### Multiple sequence alignment

AQP2 orthologs from 8 *Plasmodium* species were identified and amino acid sequences retrieved using PlasmoDB: PF3D7_0810400, PPRFG01_0811800, PmUG01_14045900, PVP01_1429900, PKNH_1430300, PBANKA_1427100, PY17X_1429200 and PGAL8A_00525700. Multiple sequence alignments were performed with T-Coffee v11.00 (37), and the resulting alignments were presented and annotated using ESPript3.0 (38).

### Phylogenetic analysis

Pore-forming motifs were identified, and the 60 amino acids surrounding each motif were extracted for multiple sequence alignment performed in R using the msa package (v1.26). Distance matrices calculated by the JC69 method using ape (v5.6.2). Phylogenetic trees were estimated using the Neighbor-Joining method (39) and generated using the ggtree package (v3.2.1), adapted from code produced previously (40). Bootstrap values were derived from 1000 iterations using ape (v5.6.2).

### Prediction of tertiary structures using Alphafold

Alphafold Monomer v2.0 (17, 18) was used to predict protein tertiary structures. For folding of PfAPQX, sequences were first adapted manually, and tertiary structures were predicted using the ColabFold server, which uses sequence alignments generated by MMseqs2 and HHsearch followed by Alphafold Monomer v2.0 to fold custom sequences (41). 3D structures were visualized and colored in iCn3D v3.18.1 (42).

### Prediction of membrane topology

Membrane-spanning helices presented in tertiary structures produced by Alphafold and visualized in iCn3D were cross-referenced with predicted membrane-spanning sequences predicted by PPM2.0 (43). Predicted membrane topology was visualized, annotated, and presented using Protter (44).

### Mosquito maintenance

*A. coluzzii* mosquitoes of Ngousso strain were maintained in an insectary at 26°C and 70% relative humidity, with a 12-hour day/night cycle with 30 min dawn/dusk transitions, respectively.

### *P. falciparum* culturing

*Plasmodium falciparum* malaria parasites of NF54 strain were maintained in 10mL human red blood cell cultures at 5% haematocrit (HTC) in complete medium (CM: RPMI-1640-R5886, hypoxanthine (0.05g/L), L-glutamine (0.3mg/L), and 10% sterile human serum). Human serum was batch tested for compatibility with gametocyte production. Asexual parasite cultures were maintained by dilution once weekly to limit parasitaemia to <10%, and media refreshed daily. Cultures were fumigated with malaria gas mixture (5% O_2_, 5% CO_2_, and 90% N_2_) and maintained in static incubators at 37°C.

### *P. falciparum* mosquito infections

*A. coluzzii* Ngousso mosquitoes separated into pots the day prior to feeding for acclimatization were infected with *P. falciparum* by standard membrane feeding as previously described (45). Briefly, gametocytes were induced from asexual *P. falciparum* cultures by establishment of 1% parasitemia cultures in reduced conditioned media, resulting in a 5% HTC culture in 8mL volume. Cultures were maintained in reduced volume at 37°C and in malaria gas atmosphere for 14 days until mosquito feed. Parasite cultures were pelleted and resuspended with fresh RBCs (200μ per culture flask) and equal volume of human AB serum. Infected blood was loaded into membrane feeders thinly covered by Parafilm connected to a water bath at 37°C, and feeders were introduced to the mosquito pots. Feeding was enabled for 10 min before mosquitoes were removed and stored in secondary containment in an incubator at 27°C and 70% relative humidity to establish infection. Blood fed mosquitoes were kept in the dark for 48 hours post-feed with no sugar feed provided to ensure survival of only fed females. After this period, mosquitoes were provided with 10% sucrose until dissection.

### Generation of AQP2-dis-pDC2 plasmid for disruption of PfAQP2

The pDC2 plasmid kindly provided by Marcus Lee (Sanger Institute) was digested and linearized with BbsI (New England Biosciences) to create TATT/AAAC overhangs downstream of the U6 promoter for insertion of the gRNA sequence. The gRNA-encoding DNA oligomers (410-432 bp of PfAQP2 coding sequence) with appended TATT/AAAC (**Table S5**) were annealed and phosphorylated with T4 polynucleotide kinase (PNK) enzyme prior to ligation into the linearized plasmid. Following gRNA insertion, the plasmid was prepared for insertion of donor DNA sequences by Gibson Assembly. All PCR reactions described were performed with CloneAmp HiFi polymerase (Takara) and the specified primers in an Applied Biosystems Thermocycler according to the manufacturer’s instructions. Firstly, the hDHFR resistance cassette was cloned from pDC2 using the P5/P6 primer pair (**Table S5**). Homology arms of 863bp upstream and 593bp downstream of the gRNA target site were cloned from *P. falciparum* genomic DNA using primer pairs P1/P2 and P3/P4, respectively, incorporating overlap sequences for Gibson Assembly into the PCR product. The pDC2 backbone was linearized by restriction enzyme digest with ApaI/AatII (New England Biosciences). DNA fragments were assembled using the NEBuilder kit (New England Biosciences) according to the manufacturer’s instructions, reconstituting the pDC2 plasmid with AQP2 5’ and 3’ homology regions flanking hDHFR.

### Transfection of *P. falciparum* by electroporation

Plasmids constructed as described above were produced in *E. coli* by transformation of constructs into competent cells followed by liquid culture of clonal colonies; plasmids were then isolated using an endotoxin-free QIAGEN maxiprep kit. 50μg of each plasmid per transfection was ethanol precipitated overnight (2.5 volumes 100% ethanol, 1/10 volumes sodium acetate (3M, pH5.2), - 20°C). Pellets were washed in 70% ethanol and resuspended in sterile TE buffer (QIAGEN). 50μg of fully dissolved plasmid was taken into a final volume of 50μL sterile TE and added to 350μL sterile incomplete cytomix (120mM KCl, 0.15mM CaCl_2_, 2mM EGTA, 5mM MgCl_2_, 10mM K_2_HPO_4_, 10mM KH_2_PO_4_, 25mM HEPES pH7.6). 5mL *P. falciparum* cultures at 5% hematocrit were synchronized with 5% sorbitol in advance of transfection; cultures were established to achieve 5-6% early ring stage parasitemia on the day of transfection. Cultures were centrifuged and washed with cold incomplete cytomix, then resuspended in complete cytomix (containing either 50μg pDC2-dis-AQP2 and 400μL transferred into an electroporation cuvette. Parasites were electroporated in a BioRad GenePulser at 310V and 950μF, with time constant in the range 7-12ms. Parasites were washed by transfer into 10mL chilled culture medium and pelleted; supernatant containing debris and cytomix was removed. Transformed parasites were transferred into a 10mL culture (3% hematocrit) and left to recover for 24 hours before drug treatment. For pDC2-dis-AQP2 transfectants, 2.5nM WR99210 was applied for 5 days. Cultures were assessed by thin smear after 2-3 weeks; once parasitemia became detectable by thin smear, clones were isolated by limiting dilution

### Isolation and genotyping of transfectant *P. falciparum* by limiting dilution

Isolation of individual transfectant clones was performed by adaptation of a limited dilution protocol. Once parasitemia of transfectant cultures exceeded 4%, thin blood smears were taken to accurately count parasitemia. RBCs in the culture were enumerated in a hemocytometer to calculate the number of infected red blood cells/mL. Once the concentration of parasites in the culture was known, a tube containing exactly 10^6^ parasites/mL in complete culture medium was established, from which serial dilutions were made to prepare 10^4^ parasites/mL in complete culture medium at 1% hematocrit (HTC). From this culture, two successive 10X dilutions followed by six successive 2X dilutions into complete medium, 1% HTC, were performed, generating parasite concentrations of 1000, 100, 50, 25, 12.5, 6.25, 3.125, and 1.5625 parasites/mL. A 96-well plate was prepared, and 100μL of each dilution was pipetted into each well of successive rows; this resulted in 100, 10, 5, 2.5, 1.25, 0.625, 0.3125, and 0.15625 parasites per well of each row A-H, respectively. The 96-well plate was maintained inside a microisolator containing malaria gas mixture. Media was changed three times per week, and HTC increased from 1% to 5% by introducing 1μL fresh RBCs on days 4, 10, 16, and 19 after establishing the plate. Rows A-B were used to establish success of parasite growth on day 10 post-smear. If parasites were detectable by thick blood smear in control wells A/B, then on day 21 clonal rows F, G, and H were examined by thick blood smear to detect parasites. Rows F-H should all contain <1 parasite per well; rows with <35% positive wells were therefore considered successful, and clones were taken into static culture flasks to recover. Once clones were at sufficiently high parasitemia, a subset of the culture was taken for gDNA extraction followed by genotyping by PCR (with remaining parasites maintained in culture pending success of genotyping). Genotyping PCR reactions were performed on 10ng of isolated parasite DNA using CloneAmp HiFi polymerase premix (Takara) in an Applied Biosystems thermocycler according to manufacturer’s instructions.

### *P. falciparum* oocyst enumeration

*A. coluzzii* mosquitoes infected with *P. falciparum* were killed on the date of dissection with CO_2_, followed by immersion in 70% ethanol. Carcasses were dissected in PBS under dissecting microscopes, unless at timepoints exploring sporozoite development, in which samples were dissected in 5% sodium azide. Following dissection, mosquito midguts were stained with mercurochrome for oocyst enumeration. Briefly, midguts were immersed in 0.1% mercurochrome for 20 minutes, prior to fixing in 4% formaldehyde for 30 minutes. Guts were washed in PBS and mounted in glycerol for analysis by light microscopy.

### Generation of *P. berghei Δpbaqp2* and *pbaqp2::3xha*

We used the *PlasmoGEM* vectors PbGEM_321755 and PbGEM-066039 (46) for PbAQP2 disruption and C-terminal 3xHA tagging, respectively. These vectors contain the human DHFR (hDHFR) selection cassette that confers resistance to pyrimethamine and the negative selection cassette yFCU that substitutes delivery of 5-fluorocytosine (47). The targeting cassettes were released by NotI digestion resulting in 74% deletion of PbAQP2 coding sequence at the gene 5’ end. Transfection, drug selection of transgenic parasites and clonal selection by dilution cloning was carried out in the *c507* line as described (20).

### Parasite genotypic analysis

*P. berghei* genomic DNA was extracted from blood sampled from positive mice using the DNeasy kit (QIAGEN) following the manufacturer’s instructions. *P. falciparum* parasite cultures at 8-10% parasitaemia were pelleted and washed with cold PBS. Cells were resuspended in cold saponin (0.05% in PBS) lysis buffer and left on ice for 5 minutes. Lysate was centrifuged (6000rcf, 5 minutes) and genomic DNA (gDNA) extracted from pellets also using the DNeasy kit (QIAGEN). Successful gene modification events or maintenance of the wildtype loci was performed by PCR using primers listed in **Table S5**.

### Exflagellation assays

Blood from *P. berghei* infected mice exhibiting 1-2% gametocytemia or from *P. falciparum* gametocyte cultures was added to RPMI medium (RPMI 1640, 20% v/v FBS, 100 μM xanthurenic acid, pH 7.4) in a 1:40 ratio and incubated for 10 min. Exflagellation events were counted in a standard hemocytometer under a light microscope.

### *P. berghei* macrogamete to ookinete conversion assays

Blood from mice exhibiting 1-2% gametocytemia was added to RPMI medium (RPMI 1640, 20% v/v FBS, 100 μM xanthurenic acid, pH 7.4) and incubated for 24 hours at 21°C to allow for ookinete formation. This suspension was then incubated with a Cy3-labelled 13.1 mouse monoclonal α-P28 (1:50 dilution) for 20 min on ice. The conversion rate was calculated as the percentage of Cy3 positive ookinetes to Cy3 positive macrogametes and ookinetes.

### *P. berghei* mosquito infections

*A. coluzzii* Ngousso mosquitoes were infected by direct feeding on mice infected with *P. berghei* at a parasitemia and gametocytemia of 5-6% and 1-2%, respectively. To determine oocyst load, midguts were dissected at 7-10 days pbf and fixed in 4% paraformaldehyde. To determine sporozoite load, 25-30 midguts and salivary glands were dissected 15 and 21 days pbf, respectively and homogenized, and sporozoites were counted in a standard hemocytometer under a light microscope. To assess mosquito-to-mouse transmission, about 30 *A. coluzzii* mosquitoes that had blood-fed on *P. berghei*-infected mice 20-22 days earlier were allowed to feed on 2-3 anaesthetized C57/BL6 mice. Mouse parasitemia was monitored until 14 days post mosquito bite by Giemsa staining.

### Western blot analysis

To extract whole cell lysates, purified blood stages, gametocytes and ookinetes were suspended in whole cell lysis buffer (1XPBS, 1% v/v Triton X-100). Proteins resolved by SDS-PAGE were transferred to a polyvinylidene difluoride (PVDF) membrane. Detection was performed using rabbit α-HA (Cell Signaling Technology) (1:1000), goat α-GFP (Rockland chemicals) (1:1000) and 13.1 mouse monoclonal α-P28 (1:1000) antibodies. Secondary horseradish peroxidase (HRP) conjugated goat α-rabbit IgG, goat α-mouse IgG antibodies (Promega) and donkey α-goat IgG (Abcam) were used at 1: 2,500, 1: 2,500 and 1: 5,000 dilutions, respectively. All primary and secondary antibodies were diluted in 5% w/v milk-PBS-Tween (0.05% v/v) blocking buffer.

### Indirect immunofluorescence assays

*P. berghei* blood stage gametocytes, ookinetes and sporozoites were fixed in 4% paraformaldehyde (PFA) in PBS for 10 min at room temperature. Fixed parasites were washed 3X with 1XPBS for 10 min each and then smeared on glass slides. Permeabilization of the parasites was done using 0.2% v/v Triton X-100 in PBS for 10 min at room temperature. Permeabilized parasites were washed 3 times in PBS for 10 min each and blocked with 1% w/v bovine serum albumin in PBS for 1 hour at room temperature. Parasites were stained with rabbit α-HA (CST) (1:1000), rabbit α-GFP (ThermoFisher) (1:500), 13.1 mouse monoclonal α-P28 (Ref) (1:1000) and 2A10 mouse monoclonal α-PfCSP (1:100) antibodies. Alexa Fluor rabbit 488 and mouse 568 conjugated secondary goat antibodies (ThermoFisher) were used at a dilution of 1:1000. 4’,6-diamidino-2-phenylindole (DAPI) was used to stain nuclear DNA. Images were acquired using a Leica SP5 MP confocal laser-scanning microscope and visualized using Image J.

## Statistical analysis

All statistical analyses were performed using GraphPad Prism v8.0.

## Acknowledgements

We thank Temesgen Menberu Kebede for outstanding assistance with mosquito and *P. berghei* culturing in mice and Maria Grazia Inghilterra for *P. falciparum* culturing. We are grateful to Marcus Lee for sharing pDC2 plasmid (Sanger Institute) and to Chris J. Janse and Blandine Franke-Fayard (Leiden University Medical Center) for sharing *P. falciparum* transfection protocols and relevant discussions.

## Author contributions

A.J.B. performed research, analyzed data, and provided draft text; C.V.U. performed research, validated results, and analyzed data; M.G. performed research; T.R.B.B. performed research; G.K.C. conceived the study, acquired funding, and wrote the paper; D.V. conceived the study, designed research, analyzed data, acquired funding, and wrote the paper.

## Funding

The work was funded by a Wellcome Trust Investigator Award to G.K.C. (107983/Z/15/Z) and a Medical Research Council (MRC) project grant (MR/T000929/1) to G.K.C. and D.V. A.J.B. was supported by a Wellcome Trust PhD fellowship award to Imperial College London (102126/B/13/Z).

## Declaration of interests

The authors declare no competing interests.

## Inclusion and diversity statement

We support inclusive, diverse, and equitable conduct of research.

## Notes

### Competing Interest Statement

The authors have declared no competing interest.

