## Supplementary Material for "Intracellular *Plasmodium* aquaporin 2 is required for sporozoite production in the mosquito vector and malaria transmission"

### Supplementary figures

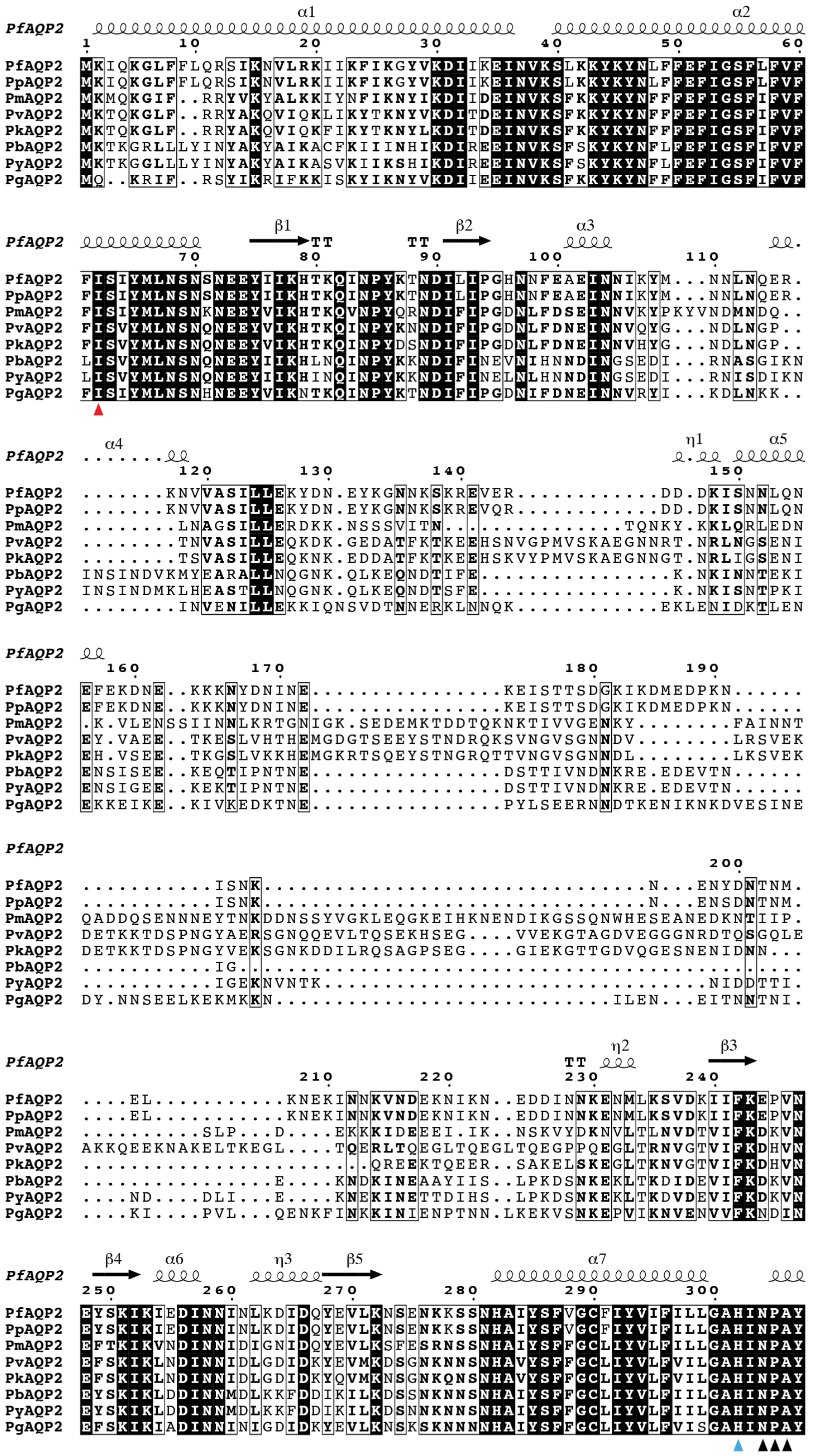

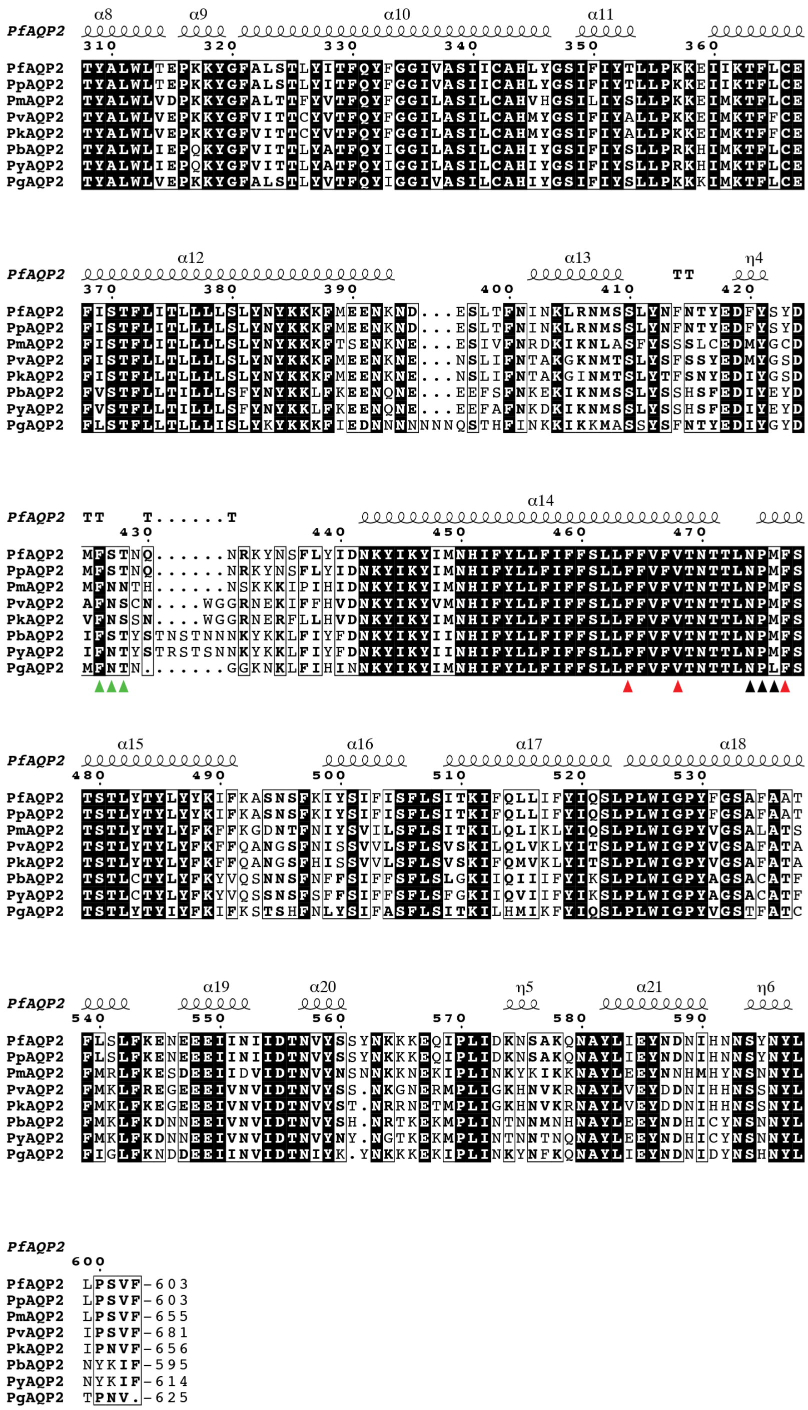

**Fig. S1. Multiple sequence alignment of *Plasmodium* AQP2 orthologs.** AQP2s from eight *Plasmodium* species are included: *P. falciparum* (PfAQP2, PF3D7_0810400), *P. praefalciparum* (PpAQP2, PPRFG01_0811800), *P. malariae* (PmAQP2, PmUG01_14045900), *P. vivax* (PvAQP2, PVP01_1429900), P. *knowlesi* (PkAQP2, PKNH_1430300), *P. berghei* (PbAQP2, PBANKA_1427100), *P. yoelli* (PyAQP2, PY17X_1429200) and *P. gallinaceum* (PgAQP2, PGAL8A_00525700). Multiple sequence alignment is performed with t-coffee, while secondary structure predictions of PfAQP2 were obtained from Alphafold Monomer v2.0 predictions, annotated using ESPript3.0 and presented above the alignment. The selectivity filter residues are highlighted with red triangles and the histidine residue participating in the pore with a cyan triangle.

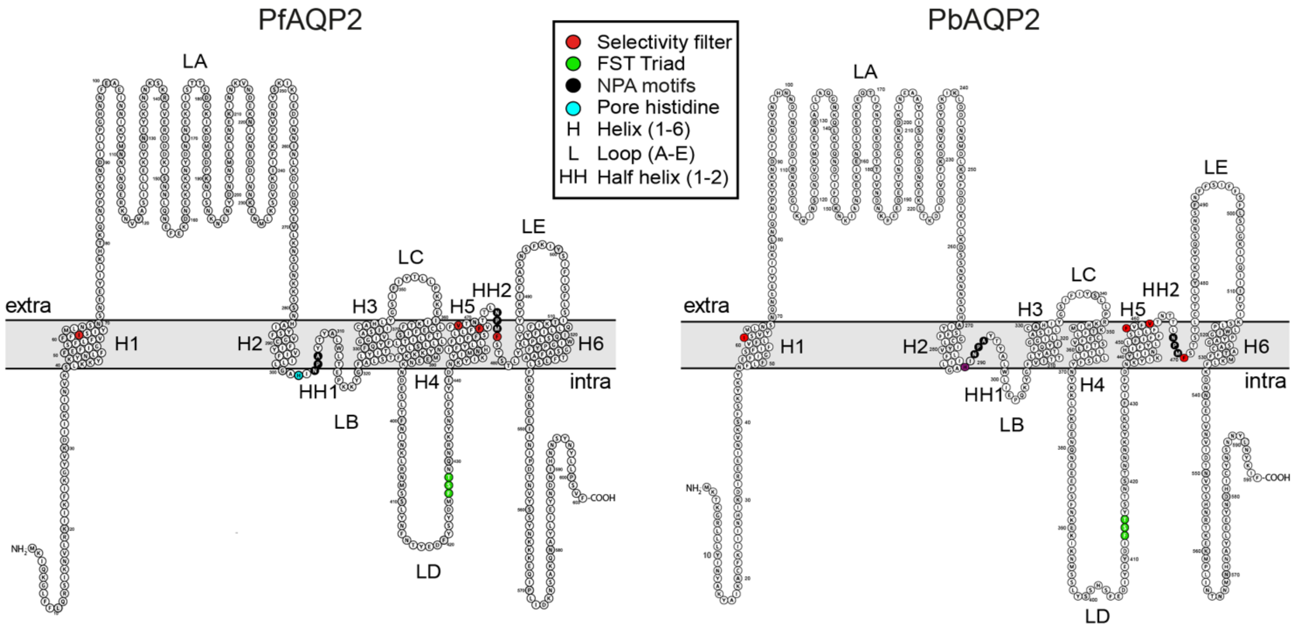


**Fig. S2 Predicted membrane topology of PfAQP2 (left) and PbAQP2 (right).** Topology of is predicted with Alphafold and PPM, and visualized using Protter. Transmembrane helices are labeled as H1-6, half-helices as HH1-2, and extramembrane loops as LA-E. Key residues are highlighted: selectivity filter in red, NPA/NPM motifs in black, FST triad of loop D in green, and histidine participating in the pore in cyan.

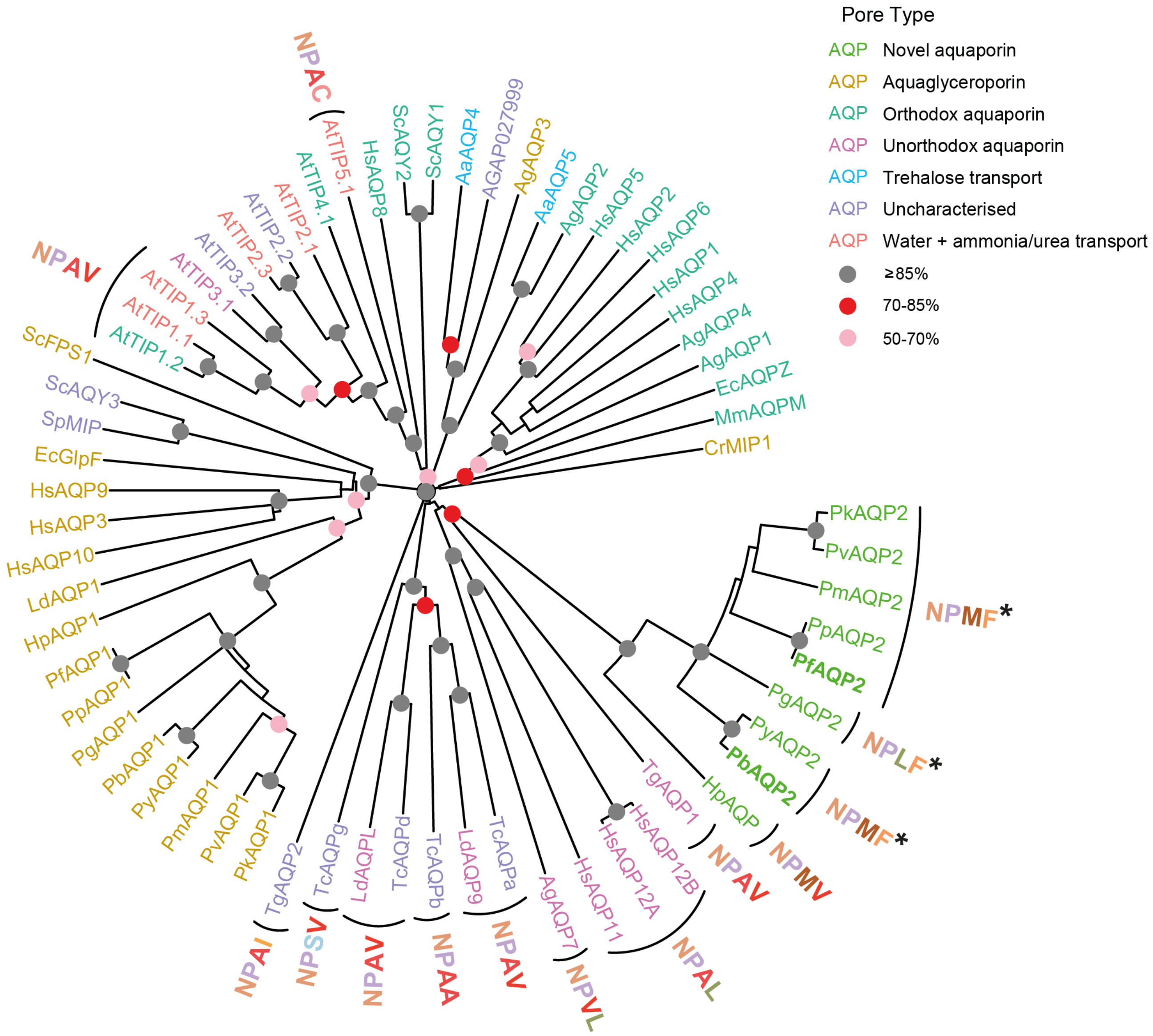


**Fig. S3. Phylogenetic tree of the conserved aquaporin pore-forming regions.** 66 aquaporin genes from 21 species are presented: *P. falciparum* (Pf), *P. berghei* (Pb), *P. praefalciparum* (Pp), *P. malariae* (Pm), *P. vivax* (Pv), *P. knowlesi* (Pk), *P. yoelli* (Py), *P. gallinaceum* (Pg), *Hepatocystis sp.* (Hp), *Toxoplasma gondii* (Tg), *Trypanosoma cruzii* (Tc), *Leishmania donovani* (Ld), *Homo sapiens* (Hs), *Anopheles gambiae* (Ag), *Aedes aegypti* (Aa), *Arabidopsis thaliana* (At), *Saccharomyces cerevisiae* (Sc), *Schizosaccharomyces pombe* (Sp), *Escherichia coli* (Ec), *Methanothermobacter marburgensis* (Mm), *Chlamydomonas reinhardtii* (Cr). Ar/R selectivity filter motifs with arginine substitutions are annotated (canonical sequence: NPAR), colored by amino acid sequence. The *Plasmodium* AQP2 genes with unique phenylalanine substitution are highlighted with an asterisk. Gene names are coloured by category of pore type and nodes are labeled with bootstrap confidence values derived from 1000 iterations as indicated in the key.

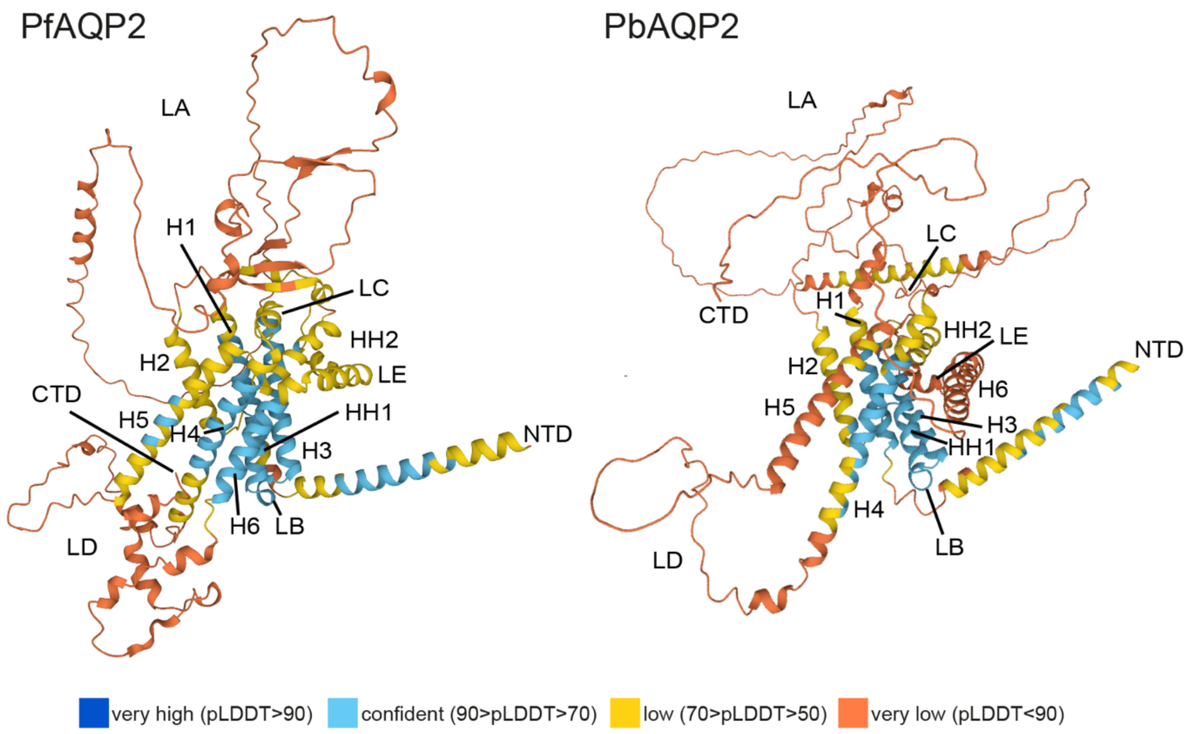


**Fig. S4. Putative 3D structures of PfAQP2 and PbAQP2.** Protein structures are predicted by Alphafold Monomer v2.0 and can be accessed in UniProt, PlasmoDB and other databases as C0H4T7 and A0A509AUY5, respectively. Structures are colored by Alphafold prediction confidence values as shown in the key below the structures.

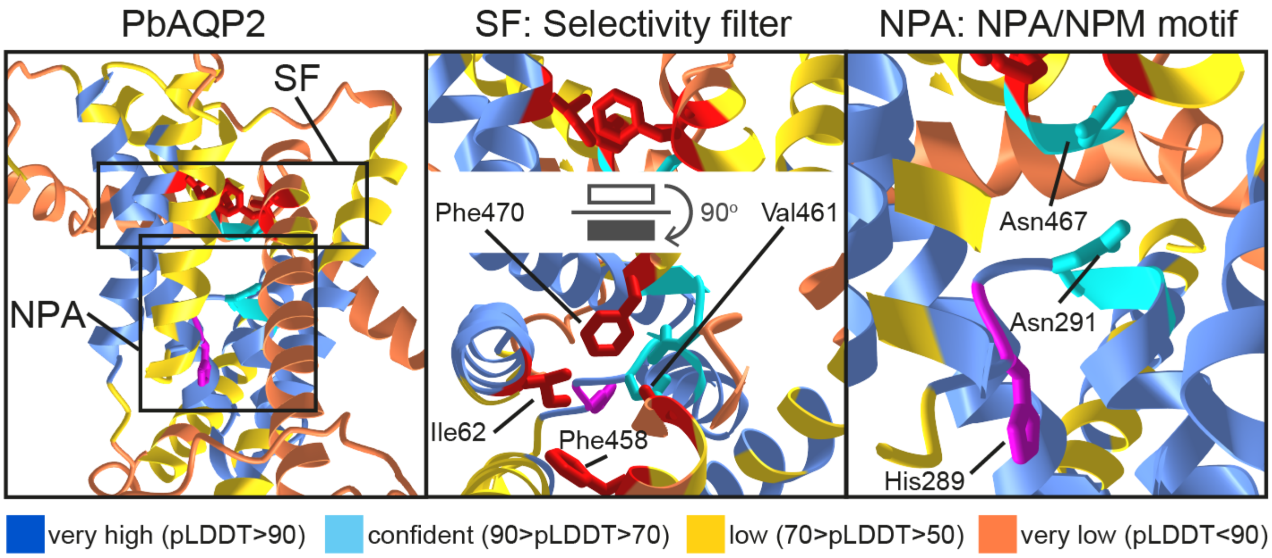


**Fig. S5. Predicted structure of PbAQP2 pore and selectivity filter.** Structures are predicted with Alphafold Monomer v2.0 on the CoLab server. Top ranking models are shown. Structures are colored by confidence level as shown in the key below the figure. Residues contributing to the selectivity filter are shown in red and those contributing to the NPA/NPM motif are shown in cyan, while the histidine residue participating in the pore is shown in magenta.

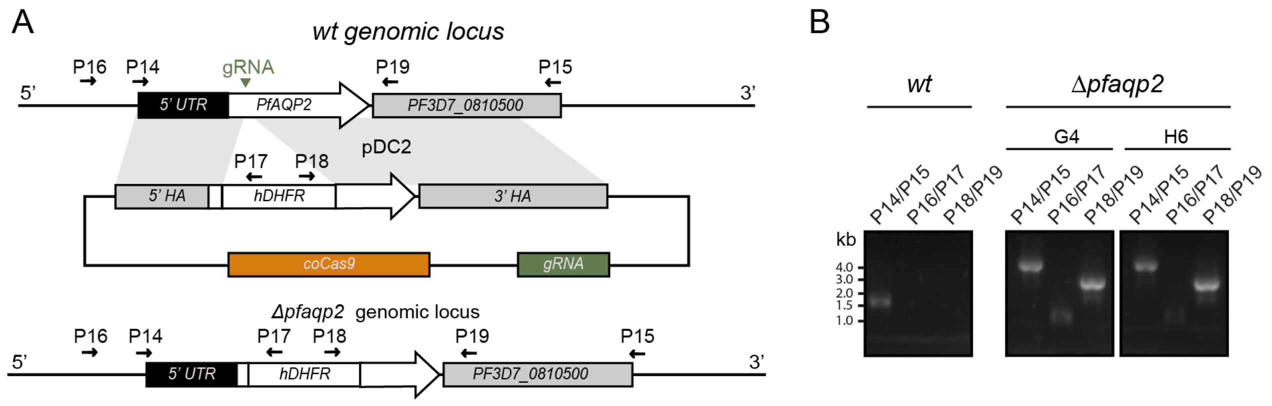


**Fig. S6. Schematic diagram of *Δpfaqp2* knockout generation by CRISPR-Cas9.** (*A*) The endogenous *NF54* PfAQP2 locus shown at the top was disrupted using the AQP2-dis-pDC2 plasmid construct shown in the center of the panel resulting in the disrupted locus shown at the bottom. For the generation of the plasmid construct, primers described in Table S5 are used to clone the 5′ and 3′ donor homology arms (HA) upstream and downstream of the Cas9 cut site on the parasite genome, respectively. The donor AQP2-dis-pDC2 plasmid includes the hDHFR selectable marker cassette. (*B*) PCR genotypic analysis of successfully generated recombinants (left picture) and isolated *Δpfaqp2* clonal populations (right picture) using primers specific for recombinant genomic sequence.

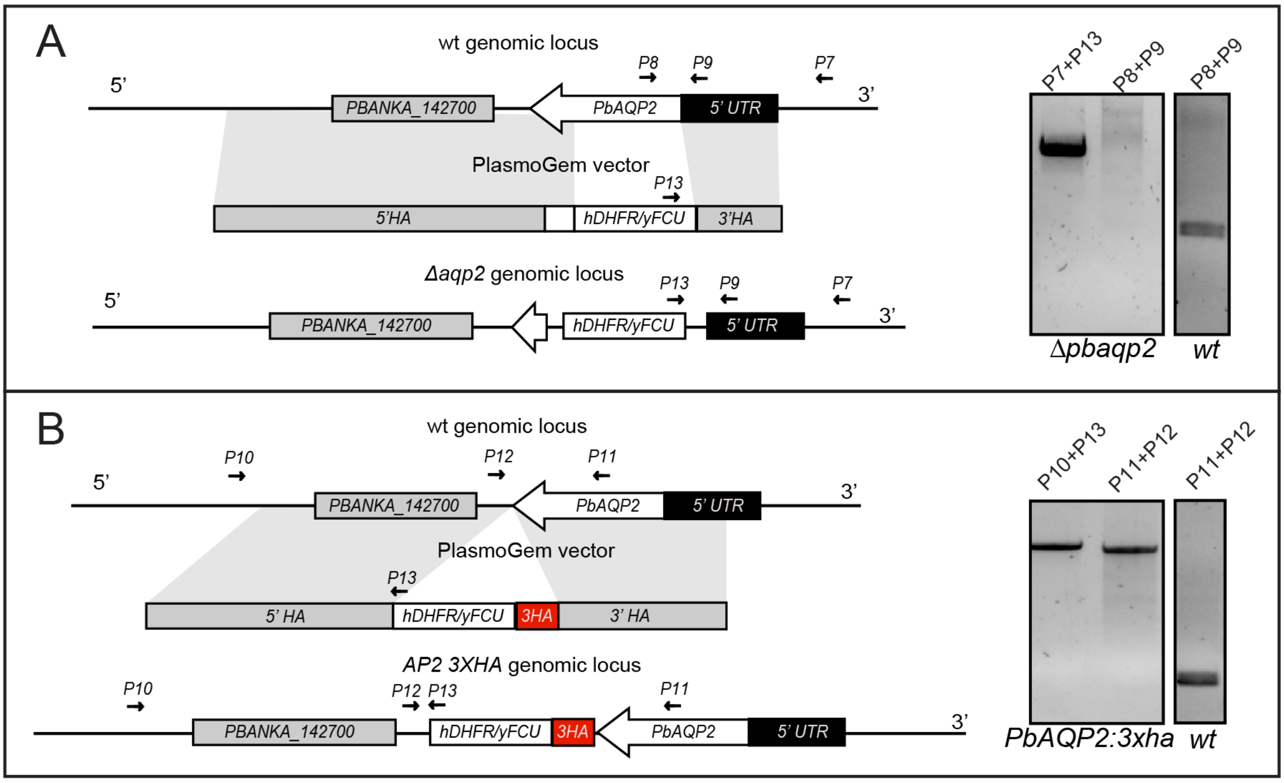


**Fig. S7. Generation of *Δpbaqp2* mutant and gene tagged parasites.** (*A*) Schematic representation of the disruption of *P. berghei* in the *c507* line, using *Plasmo*GEM vectors carrying the *human DHFR* (*hDHFR)/yFCU* pyrimethamine resistance cassette. In each panel, the *wt* genomic locus (top), the transfected DNA fragment carrying the homologous arms (HA) and the selectable markers (middle), and the final transgenic locus after gene disruption by double crossover homologous recombination (bottom) are shown. Constructs are not drawn to scale. Small arrows show the primers used for the confirmation of integration of gene targeting constructs and deletion of the gene of interest using PCR as shown in the panels on the right. The *c507 wt* parental parasites were used as a control for detection of the *wt* locus. (*B*) Schematic representation of the 3XHA tagging of in the c507 line by using PlasmoGEM gene tagging vectors that carry the hDHFR)/yFCU selectable marker. In each panel, the wt genomic locus (top), the plasmid carrying the homologous arms (HA), the transfected DNA fragment carrying the tagging cassettes and the selectable markers (middle), and the final transgenic locus after double crossover homologous recombination (bottom) are shown. Constructs are not drawn to scale. Small arrows show the primers used for the confirmation of integration of gene targeting constructs and deletion of the gene of interest using diagnostic PCR reactions as shown in the panels on the right**.**

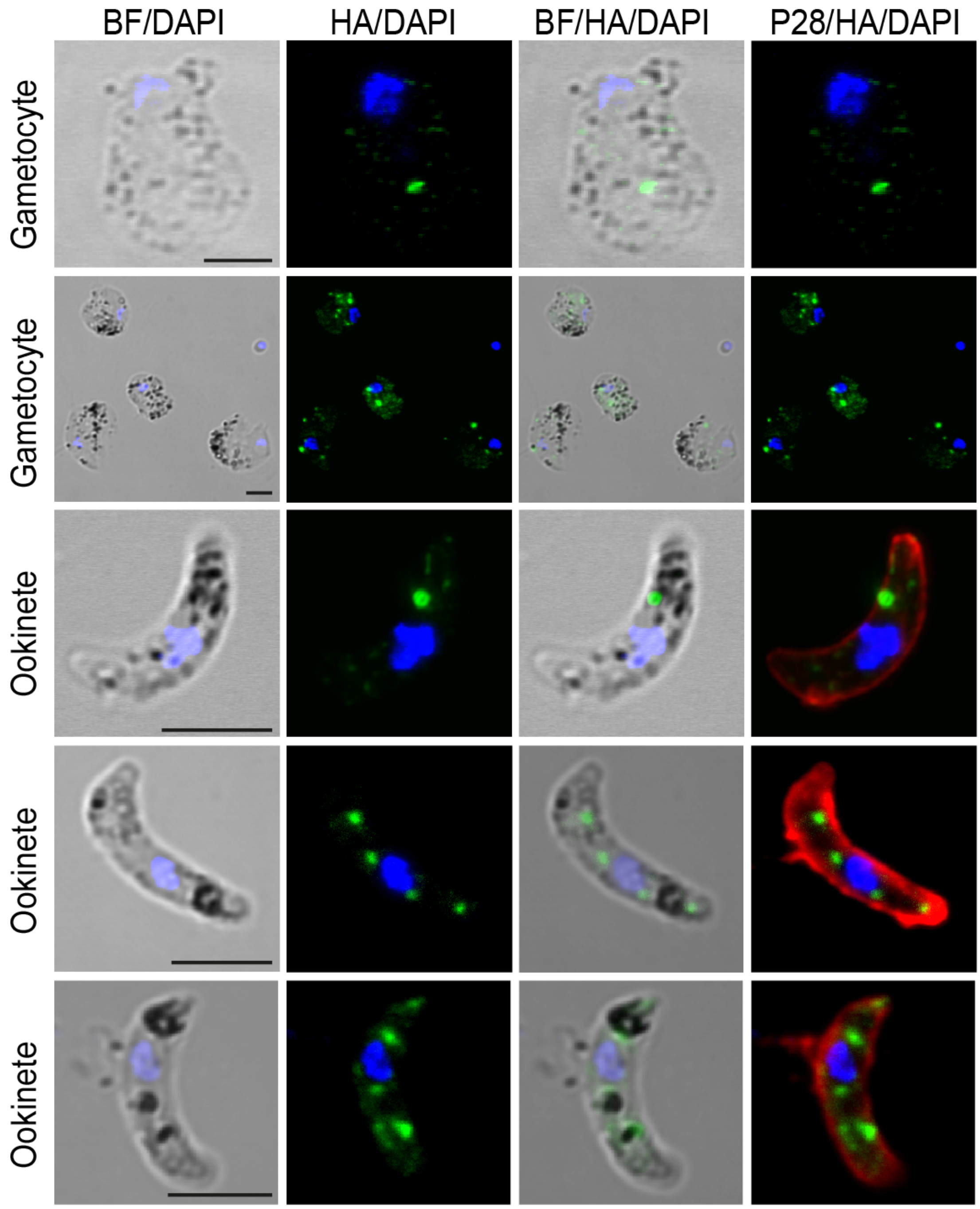


**Fig. S8. Subcellular localization of *P. berghei* AQP2.** Immunofluorescence assays of *aqp2::3xha* purified gametocytes, *in vitro* cultured ookinetes sampled 24 hours after culture setup, and sporozoites obtained from infected *A. coluzzii* midguts 9 days pbf. AQP2::3xHA (green) and P28 (red, ookinetes) or CSP (red, sporozoites) are detected using antibodies. DNA is stained with DAPI (blue). BF is brightfield. Scale bar is 2.5 μM.

#### Supplementary tables

**Table S1.** Oocyst counts of control *Δpfaqp2* mutant parasites and NF54 controls in *A. coluzzii*

| Replicar | Parasite | No of midguts | Prevalence (%) | Arithmetic mean | Median | Parasite Range | *P* value |
|---|---|---|---|---|---|---|---|
| Pool | *NF54* | 139 | 69.8 | 2.25 | 1 | 28 | |
| | *G4* | 110 | 65.5 | 1.74 | 1 | 15 | 0.7988 |
| | *H6* | 141 | 68.1 | 1.82 | 1 | 24 | 0.8858 |
| R1 | *NF54* | 22 | 54.5 | 0.77 | 1 | 3 | |
| | G4 | 22 | 50.0 | 0.86 | 0.5 | 5 | >0.9999 |
| | H6 | 25 | 72.0 | 1.44 | 1 | 6 | 0.2312 |
| R2 | *NF54* | 21 | 57.1 | 1.91 | 1 | 12 | |
| | G4 | 18 | 55.6 | 0.78 | 1 | 3 | 0.6934 |
| | H6 | 27 | 55.6 | 0.96 | 1 | 3 | >0.9999 |
| R3 | *NF54* | 61 | 75.4 | 2.95 | 2 | 28 | |
| | G4 | 30 | 73.3 | 2.23 | 2 | 7 | >0.9999 |
| | H6 | 44 | 70.5 | 2.39 | 1.5 | 24 | 0.6039 |
| R4 | *NF54* | 35 | 77.1 | 2.17 | 1 | 11 | |
| | G4 | 40 | 72.5 | 2.28 | 2 | 15 | >0.9999 |
| | H6 | 45 | 71.1 | 2.00 | 2 | 10 | >0.9999 |

Infection data from four biological replicates of *A. coluzzii* infected with *Δpfaqp2* (clones G4 and H6) *and NF54* parasite lines, assessed at 9 days post-infection. *P* values were calculated using the Kruskal-Wallis test, comparing to wild type, and adjusted with Dunn's multiple comparisons test.

**Table S2.** Sporozoite numbers of control *NF54* and *Δpfaqp2* parasites in *A. coluzzii* infections

| | Midgut sporozoites | | | Salivary gland sporozoites | | |
|---|---|---|---|---|---|---|
| Parasite | Mean | SEM | P value | Mean | SEM | P value |
| *NF54* | 3717 | 967.1 | | 1149 | 211.0 | |
| G4 | 30.4 | 8.7 | < 0.0001 | 28.5 | 13.5 | < 0.0001 |
| H6 | 30.5 | 16.7 | < 0.0001 | 20 | 6.6 | < 0.0001 |

Mean oocyst and salivary gland sporozoite numbers of *A. coluzzii* infected with *Δpfaqp2* (clones G4 and H6) *and NF54* parasite lines. Pooled data from 4 biological replicates are presented. In each replicate, sporozoite numbers were determined from 25-30 homogenized mosquito midguts or salivary glands at 11 and 18 days pbf, respectively. *P* values were calculated by one-way ANOVA, comparing to wild type, and adjusted with Dunnett's multiple comparisons test. SEM represents standard error of mean.

**Table S3.** Oocyst numbers of control *c507* and *Δpbaqp2* parasites in *A. coluzzii* infections

| Replicate | Parasite | No. of midguts | Prevalence (%) | Mean | Median | Range | *P* value |
|---|---|---|---|---|---|---|---|
| Pool | *c507* | 61 | 82 | 22.1 | 13 | 0-176 | |
| | *Δpbaqp2* | 64 | 78 | 16.4 | 9 | 0-185 | 0.2284 |
| R1 | *c507* | 32 | 81 | 22.8 | 13 | 0-176 | |
| | *Δpbaqp2* | 31 | 84 | 13.1 | 9 | 0-57 | 0.581 |
| R2 | *c507* | 29 | 83 | 21.2 | 14 | 0-114 | |
| | *Δpbaqp2* | 33 | 73 | 19.5 | 8 | 0-185 | 0.2941 |

Data are from two biological replicates of infected *A. coluzzii* dissected at 8 days pbf. *P* values were calculated using the Mann-Whitney t-test.

**Table S4.** Sporozoite numbers of control *c507* and *Δpbaqp2* parasites in *A. coluzzii* infections

| | Midgut sporozoites | | | Salivary gland sporozoites | | |
|---|---|---|---|---|---|---|
| Parasite | Mean | SEM | P value | Mean | SEM | P value |
| *c507* | 3,567 | 259 | | 1,843 | 83 | |
| *Δpbaqp2* | 199 | 58 | 0.0115 | 34 | 24 | 0.0045 |

Mean oocyst and salivary gland sporozoite numbers from *A. coluzzii* infections. Pooled data from two biological replicates are presented. In each replicate, sporozoite numbers were determined from 25-30 homogenized mosquito midguts or salivary glands at 15 and 21 days pbf, respectively. *P* values were calculated using the unpaired Student's t-test. SEM represents standard error of mean.

**Table S5. Oligonucleotide primers used in this study**

| Primer name | Sequence (5′ to 3′) | Description |
|---|---|---|
| P1 F | TATATGGGAATTTCCTTATAGGGCCCTTTAGGTTTTATTATTTTAGGG | PfAQP2 disruption upstream target |
| P2 R | CGCAGAGAAATCTAGAGGTACCGAGTATTGTTACCTTTATATTCGTTATC | PfAQP2 disruption upstream target |
| P3 F | TTACAAGCGAATTAGCTAAGCATGCGACGATGATAAAATAAGCAAT | PfAQP2 disruption downstream target |
| P4 R | TTCCCCGAAAAGTGCCACCTGACGTCATATAATACTGGCAACGATACC | PfAQP2 disruption downstream target |
| P5 F | CTCGGTACCTCTAGATTTCTC | hDHFR cassette forward primer |
| P6 R | GCATGCTTAGCTAATTCGCTTGTAA | hDHFR cassette reverse primer |
| P7: PlasmoGEM AQP2 GT F | TTGTGTTAGCTCTCTTGCCT | Pb diagnostic primer WT and KO/*c507* |
| P8: PlasmoGEM AQP2 QCR1 | TCAACAATGCCCTTGCCTCA | Pb diagnostic primer WT and KO/*c507* |
| P9: PlasmoGEM AQP2 QCR2 | TCATTCTACGCTTTATTAACTCA | Pb diagnostic primer WT/*c507* |
| P10: PlasmoGEM AQP2HA GT F | TGGGTGCGTACAATGACTAGA | Diagnostic primer tag |
| P11: PlasmoGEM AQP2HA QCR1 | TCTGCCTCTATGGATTGGGCCA | Diagnostic primer tag |
| P12: PlasmoGEM AQP2HA QCR2 | TGACAAGAAAAAGGTGGCAT | Diagnostic primer tag |
| P13: PlasmoGEM GW2 | CTTTGGTGACAGATACTAC | Diagnostic primer KO and tag |
| P14 | CTTTAGGTTTTATTATTTTAGGG | Pf diagnostic primer WT and KO |
| P15 | CATATAATACTGGCAACGATACC | Pf diagnostic primer WT and KO |
| P16 | GAATGTACTTATATATTTGTACACC | Pf diagnostic primer KO |
| P17 | AACATATGTTAAATATTTATTTCTC | Pf diagnostic primer KO |
| P18 | GTGTACTTGTTAAAATGCTATATCA | Pf diagnostic primer KO |
| P19 | GACAGTATTCGATGCTAAGAG | Pf diagnostic primer KO |
| P20 | TATTGAAAGTAAAAGGGAAGTGGAA | PfAQP2 disruption gRNA oligo F |
| P21 | AAACTTCCACTTCCCTTTTACTTTC | PfAQP2 disruption gRNA oligo R |
| P22 | AGCATTTGCCGCAACTTTCC | PfAQP2 qPCR primer F |
| P23 | GCGCTGTTTTTATCTATTAACGGT | PfAQP2 qPCR primer R |
| P24 | GAACCACCAAAGAGACCTTAC | EIF1α qPCR primer F |
| P25 | TCTAGCTTCTTCCAAAACTTCC | EIF1α qPCR primer R |

Overlap sequences for Gibson assembly and restriction sites are shown as underlined. F, forward; R, reverse; WT, wild-type; KO, knockout. All primers are listed in a 5′ to 3′ direction.
